# Short-term microglia depletion via CSF-1R inhibition promotes functional network reorganization and motor recovery after cortical ischemia

**DOI:** 10.1101/2025.11.07.687313

**Authors:** Sara Isla Cainzos, Fanny Quandt, Malte Borggrewe, Hanna-Marie Altjohann, Tim Magnus, Jonatan Biskamp

## Abstract

**BACKGROUND:** Pharmacological options to promote long-term rehabilitation after stroke remain limited. Microglia play a complex role in post-stroke pathology, contributing both to repair and secondary injury. How short-term depletion during the subacute phase affects functional recovery remains unknown.

**METHODS:** Wild-type mice were trained in a skilled reaching task and underwent permanent distal medial cerebral artery occlusion or sham intervention. Mice received either a colony-stimulating factor 1 receptor inhibitor or vehicle treatment between days 3 and 7 post-stroke to deplete microglia. Fine motor performance was assessed behaviorally while bilateral cortical activity was recorded longitudinally through epidural electrocorticography.

**RESULTS:** Microglia depletion did not affect infarct size but resulted in near-complete restoration of fine motor function by day 7, coinciding with maximal microglial depletion. Recovery of fine motor function was accompanied by significant functional connectivity changes in bilateral sensorimotor networks, including increased beta-band connectivity in the ipsilesional motor cortex, which correlated with contralateral fine motor improvement. After microglial repopulation, cells showed altered morphology and gene expression profiles.

**CONCLUSIONS:** Transient microglial modulation during the subacute phase after stroke promotes cortical network reorganization and motor recovery, highlighting a potential time window for future translational studies.

## Introduction

In recent years, the treatment of stroke has advanced significantly. While these developments have led to more effective acute interventions aimed at minimizing immediate brain damage, current therapeutic strategies remain primarily focused on the acute phase of stroke. Pharmacological options that influence the subsequent course of the disease and support long-term recovery are still notably limited.

Increasing attention has turned to the brain’s innate immune system as a potential modulator of recovery. In particular, microglia, the resident immune cells of the central nervous system, play a pivotal role in this context, initially promoting inflammation, later contributing to tissue repair and recovery ^1^. Microglia engage in diverse interactions with neurons, continuously surveying their environment through dynamic cellular processes and actively participating in the remodeling of synaptic connections ^2^. Moreover, their ability to shape synaptic receptor composition, modulate network oscillations both in vitro and in vivo and their role in learning and working memory has been demonstrated ^3–7^.

Over the past decade, strategies aimed at depleting microglia have emerged as promising translational approaches for various neurological disorders ^8^. The survival of microglia critically depends on signaling through the colony-stimulating factor 1 receptor (CSF-1R) ^9^. The small molecule PLX5622 selectively antagonizes CSF-1R, resulting in efficient microglial depletion and upon withdrawal of the inhibitor, repopulation of the microglial compartment occurs ^9,10^.

Recent studies have demonstrated encouraging outcomes following short-term CSF-1R inhibition in models of neuronal injury. For instance, microglial renewal in a murine model led to reduced lesion-induced release of inflammatory markers, a resolution of the activated microglial phenotype, and an improvement in behavioral deficits ^11^. Similarly, transient depletion followed by repopulation of microglia after traumatic brain injury mitigated both neuropathological changes and neurological impairments ^12,13^. Notably, Barca et al.^14^ showed that short-term CSF-1R inhibition between days 3 and 7 post-ischemia induced microglial repopulation, accompanied by alterations in glial morphology, phenotype, gene expression patterns, and cellular recruitment—changes that were associated with signs of enhanced functional recovery.

Based on these findings, we hypothesize that short-term microglial depletion in the subacute phase after ischemic stroke followed by repopulation significantly impacts neuronal activity within the penumbra and promotes functional recovery. To assess this, we evaluated changes in neuronal network activity across both the acute injury phase and the subsequent recovery period using a 16-channel electrocorticography (ECoG) electrode array implanted over both hemispheres. Fine motor deficits were evaluated over time using the single-pellet reaching task (SPR). Given that the primary motor cortex (M1) plays a critical role in mediating recovery following sensory cortical strokes, we placed particular emphasis on M1 activity after distal middle cerebral artery occlusion (dMCAo), resulting in sensory cortical stroke ^15^. The compound PLX5622 that results in efficient microglial depletion was administered daily from day 3 to day 7.

Despite no differences in infarct size between groups, mice treated with PLX5622 showed a near-complete recovery of fine motor function as early as day 7, coinciding with the peak of microglial depletion. Functional recovery was paralleled by connectivity changes in bilateral sensorimotor networks, most notably a sustained increase in beta-band connectivity within the ipsilesional motor cortex and a decrease in contralesional somatosensory gamma connectivity correlated with recovery of contralateral forepaw function. Following repopulation, microglia displayed altered morphology and a distinct transcriptomic profile, while the observed behavioral improvements remained stable. With this study, we identified transient microglial depletion as a therapeutic strategy capable of cortical network reorganization and supporting post-stroke motor recovery.

## Methods

### Data availability

The data that support the findings of this study and all custom-written MATLAB codes are available from the corresponding author upon reasonable request.

### Ethics statement

C57/Bl6 male mice aged nine postnatal weeks at the start of the experiment provided by the animal facility of the University Medical Center Hamburg-Eppendorf were used in this study. Experimental protocols were approved by the Behörde für Justiz und Verbraucherschutz der Freien und Hansestadt Hamburg (approval numbers N53/2020 and N102/2023). All procedures followed the guidelines of the animal facility of the University Medical Center Hamburg-Eppendorf and complied with the Guide for the Care and Use of Laboratory Animals. Animals were housed in groups under a 12-hour dark-light cycle. After implantation of electrodes mice were housed individually for 49 days.

Details on study design and the following methods are available in the Supplemental Material.

### Study design

To assess microglial depletion, repopulation, morphology and stroke volume a cohort of 24 wild-type mice underwent dMCAo surgery and received either PLX5622 or vehicle treatment from days 3 to 7 post-stroke. Flow cytometry (FACS) was performed on the contralesional hemisphere at days 7, 14, 21, and 28 post-ischemia (pi). Histological analysis on the ipsilesional hemisphere was performed at day 14 pi. Stroke volume was assessed at days 1, 7, 14, 21 and 28 pi with MRI (Fig. 1A).

**Figure 1.**
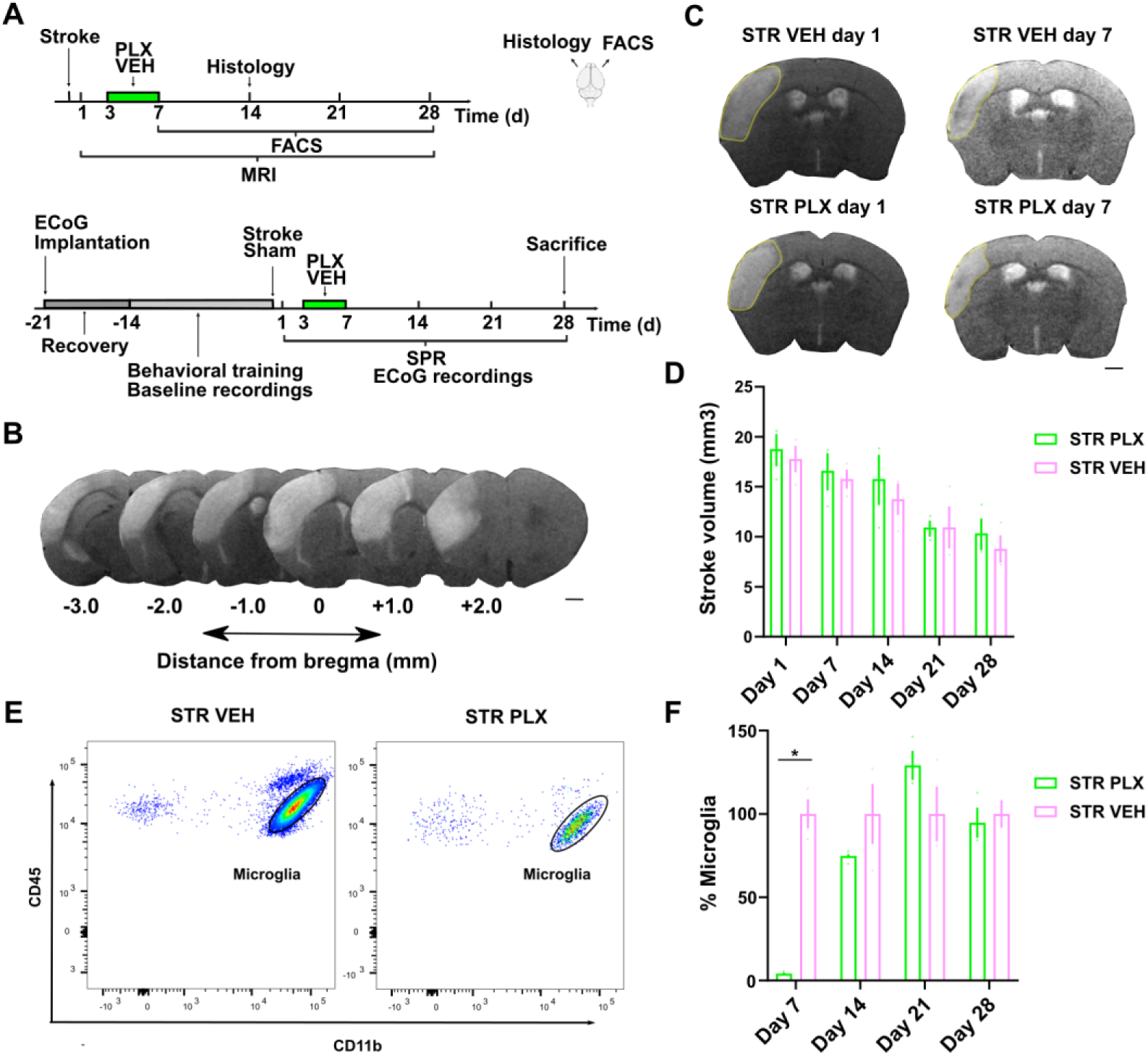
Short-term PLX5622 treatment induces microglia depletion after stroke, followed by repopulation upon drug withdrawal. (**A**) Timelines of the experiment. (**B**) Representative coronal brain MRI sections after stroke. (**C**) Representative illustration of stroke volume differences between PLX and VEH groups; lesion area outlined in yellow. (**D**) Stroke volume quantification. STR PLX: n=3; STR VEH: n=3. (**E**) Gating strategy used to assess microglial depletion. (**F**) Percentage of microglia remaining after treatment. STR PLX: n=3; STR VEH: n=3. Black scale bars in B and C indicate 1 mm. *p<0.05. Data are shown ± SEM. SH, sham; STR, stroke; PLX, PLX5622; VEH, vehicle.

To investigate the effects of microglia depletion and repopulation on functional deficits and recovery after stroke a cohort of 49 wild-type mice were implanted with ECoG electrodes and trained in a SPR task. After dMCAo or sham surgery mice received daily PLX5622 or vehicle treatment from day 3 to day 7 pi. Mice were functionally tested and recorded at days 1, 3, 7, 14, 21, and 28 pi. Before any behavioral training or electrophysiological recording started mice were granted a recovery period of one week after electrode implantation (Fig. 1A). To study the effects of microglia depletion and repopulation at a phenotype level, a third cohort of 8 wild-type mice underwent dMCAo surgery and received either PLX5622 or vehicle treatment from days 3 to 7 pi. Single-cell RNA sequencing analysis was performed at day 14 pi on pooled samples within each treatment group.

### Electrode implantation and dMCAo surgery

Electrode implantation and dMCAo surgery were performed as previously described ^15^. In short, 16 platinum-iridium electrodes of 127 μm diameter were chronically implanted epidurally covering both hemispheres, then an ischemic stroke was induced contralateral to the dominant forepaw or randomly on either hemisphere if no paw preference could be determined. As previously shown ischemic lesions were located exclusively within the sensory cortex (Fig. 1B, C) ^15^.

### PLX5622 or vehicle treatment

PLX5622 or vehicle treatment was administered orally *ad libitum* for four consecutive days, starting on day 3 after dMCAo or sham surgery. Detailed preparation and dosing information is provided in Supplemental Material.

### Infarct volume analysis using MRI

Infarct volume was assessed using a 7-Tesla small animal MR imaging system (ClinScan, Bruker, Ettlingen, Germany) with a T2-weighted MRI protocol. Infarct volumes were quantified using NIH ImageJ software (version 1.53e). Corrected stroke volumes were calculated as previously described ^16^.

### Tissue isolation, FACS, histology and microglia sorting

In short, at given time points animals were terminally anesthetized, intracardially perfused and brains were dissected.

For FACS, the contralateral hemisphere was enzymatically digested, filtered, and subjected to erythrocyte lysis and Percoll gradient centrifugation to isolate immune cells. Cells were stained targeting microglia and other immune markers, and quantitative analyses were performed to assess microglial depletion and repopulation (Fig. S1).

For histology, the ipsilesional hemisphere was coronally sectioned and microglia were labeled by immunohistochemistry. Morphology in the peri-infarct cortex was quantified by Fractal and Sholl analysis in Fiji/ImageJ2 using the FracLac plugin for fractal complexity and the Sholl Analysis plugin ^17–19^.

For single-cell sequencing, the ipsilesional cortex was dissected and enzymatically digested with papain and collagenase/DNase in the presence of transcriptional inhibitors (Actinomycin D, Triptolide, Anisomycin). The resulting cell suspension was filtered and subjected to Percoll gradient centrifugation. Microglia were stained for cell surface markers and viability, and isolated for downstream single-cell RNA sequencing.

Full experimental details are provided in the Supplemental Material.

### Single-cell sequencing analysis

Raw reads were processed with the nf-core/scRNAseq pipeline using Cell Ranger and analyzed using Scanpy ^20, 21^. Low quality cells and potential doublets were filtered, followed by normalization and selection of highly variable genes. Cells were clustered using Leiden clustering and annotated with known microglia and other myeloid markers. Differentially expressed genes in PLX5622 group versus vehicle and across distinct homeostatic subsets were determined using MAST ^22^. Gene set activity was scored using AUCell and differential activity was determined using Wilcoxon test. See Supplemental Material for detailed quality control, clustering, and annotation parameters and Tables S1-3.

### Behavior

Fine motor skills were evaluated using a SPR task as previously described ^15^. In short, mice were placed in a transparent chamber and trained to reach through a window to retrieve a food pellet placed outside. During the first week of training the dominant forepaw was determined. Then, individual baselines were defined for each mouse, averaging the number of successful reaching attempts on the last 2 days of training. Test performance was assessed in relation to individual baseline performance (in %). General locomotor activity was evaluated in an open field scenario using ANY-maze Video Tracking System 7.48 (Stoelting Co.).

### *In vivo* electrophysiology and data analysis

ECoG data were recorded from freely behaving mice (W2100 System, HS16 Headstage, Multichannel Systems) while exploring a circular arena and being videotaped ^15^. Episodes of locomotion (‘active’) as well as episodes of resting behavior with a minimum duration of 2 seconds were visually identified offline and used for electrophysiological data analysis. Data processing was performed in MATLAB using the FieldTrip toolbox ^23^. In short, continuous data were filtered (0.3 - 180 Hz), active episodes were extracted, and artifacts were manually rejected. Power spectral density (PSD) was computed on individual episodes and averaged. Peak frequencies of the PSD and aperiodic parameters were extracted in Python using the FOOOF toolbox ^15,24^. Frequency bands were defined as theta (4–10 Hz), low beta (10–20 Hz), high beta (20–30 Hz), and gamma (30–60 Hz) and relative power was calculated as the proportion of the total spectral power (1–60 Hz). Connectivity was computed as the imaginary part of the cross-spectral coherence, and spectra were averaged over time of each experiment. Group level analyses matched anatomically corresponding channels between hemispheres. For visualization, ipsilesional channels are plotted on the left. Further details of signal processing and analysis are provided in the Supplemental Material.

### Statistics

Statistics were performed in Prism (9.3.0, GraphPad) and R (4.1.1, RStudio). Data are presented as mean ± SEM and were considered statistically significant when p<0.05.

## Results

To assess the effect of short-term microglia depletion and repopulation on functional recovery and neural activity after sensory cortical stroke, mice received either the compound PLX5622 (PLX) or vehicle (VEH) treatment between day 3 and day 7 after dMCAo or sham surgery (Fig. 1A).

FACS confirmed >95% depletion of microglia by day 7 after stroke in PLX-treated animals [mean difference to VEH = -152.22, 95% CI (-273.00 to -31.44), adjusted p=0.032]. After treatment withdrawal, microglia numbers gradually repopulated (day 14, ∼75%, p=0.754; day 21, ∼130%, p=0.617; day 28; ∼100%, p=0.991; Fig. 1F).

MRI scans at day 1, 7, 14, 21, and 28 post-dMCAo revealed no significant differences in lesion volume between PLX and vehicle treated animals (Fig. 1C, D). At the group level, stroke volumes decreased from an initial ∼18 mm³ at day 1 to ∼10 mm³ by day 28. In conclusion, PLX administration between day 3 and 7 after stroke led to short-term microglia depletion and a subsequent repopulation, however, did not affect lesion volume.

Single-cell RNA sequencing of microglia cells from the ipsilateral cortex at day 14 after dMCAo revealed that repopulated microglia exhibited a distinct transcriptional profile compared with vehicle-treated samples (Fig. 2). Cells were clustered and annotated based on established marker genes (Fig. S2A) and gene set activity related to microglial functions (Fig. S2B). In the repopulated group, a population of homeostatic microglia with a more reactive phenotype (HomR) was observed. This population maintained homeostatic marker expression while upregulating reactive genes (*Apoe*, *Axl*, IFN-response) and chemokines (Fig. 2C, Fig. S2C), with increased activity of gene sets associated with disease-associated microglia (DAM), neuroprotection, and tissue repair (Fig. 2B). Within the homeostatic compartment, both the classical homeostatic cluster (Hom) and HomR showed elevated expression of genes involved in phagocytosis, tissue remodeling, and neuroprotection (*Cd36*, *Axl*, *Cd44*), with HomR additionally upregulating repair and debris clearance genes (*Grn*, *Tgm2*). Pro-inflammatory cytokines such as *Il1b* were downregulated across clusters (Fig. 2D, F), while other microglial clusters displayed enhanced expression of interferon-stimulated genes and repair-associated programs, while genes and gene sets linked to cytokine production were reduced (Fig. 2E, F).

**Figure 2.**
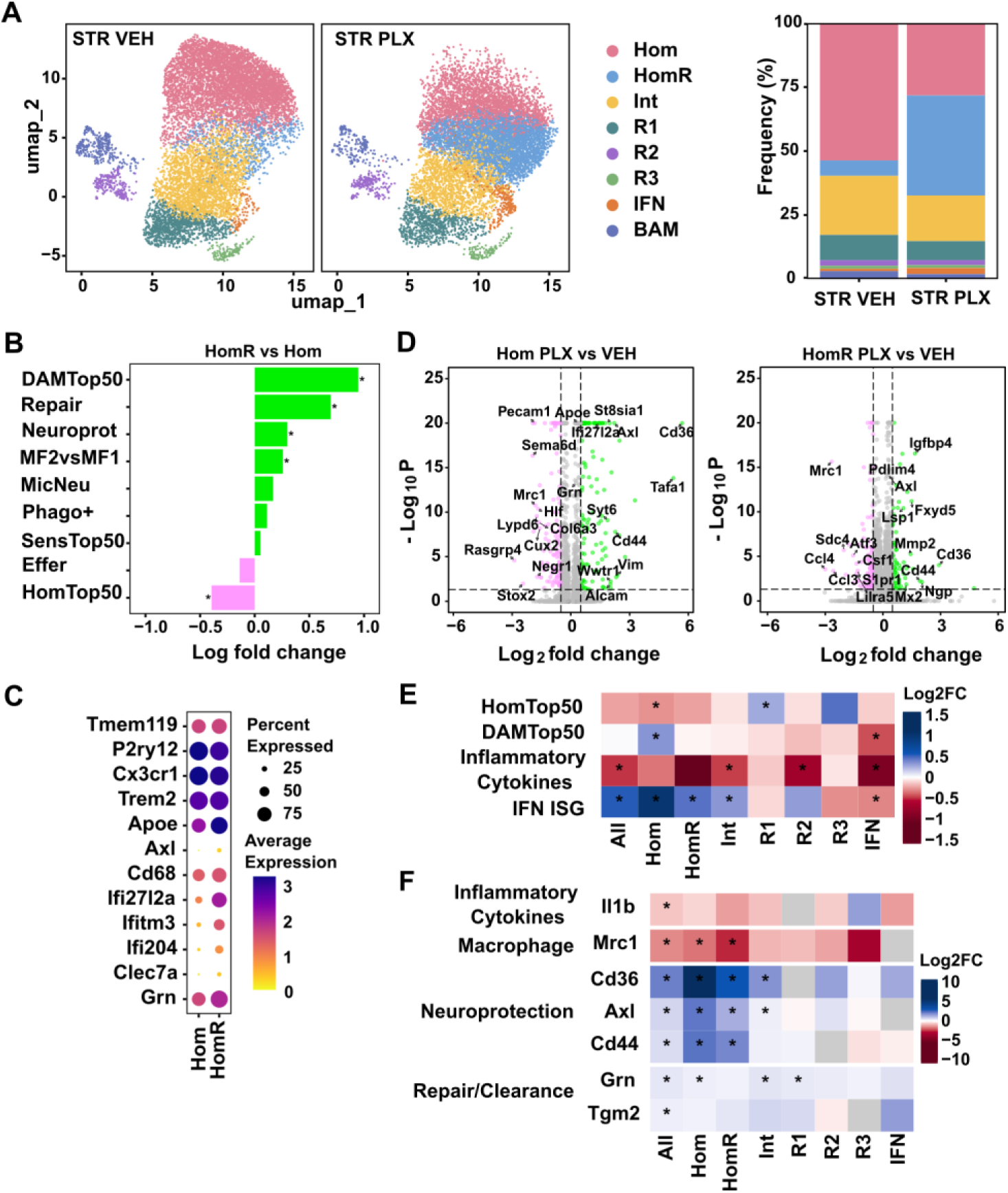
Single-cell RNA sequencing of repopulated microglia at day 14 post-stroke. (**A**) Left, separate UMAPs per condition showing microglia clustering. Right, Frecuency (%) of cells in each cluster. (**B**) Bar graph of differential GSA comparing HomR vs Hom. (**C**) Dotplot of selected genes upregulated in the PLX group. (**D**) Volcano plots of differentially expressed genes (DEGs) in selected clusters. (**E**) Heatmap of differential gene set activity (GSA) comparing PLX vs VEH. (**F**) Heatmap of selected DEGs across clusters. Cells were pooled from STR PLX: n=4 and STR VEH: n=4. STR, stroke; PLX, PLX5622; VEH, vehicle; Hom, Homeostatic; HomR, Homeostatic Reactive; Int, Intermediate; R1–R3, Reactive subtypes 1–3; IFN, IFN-responsive; Effer, efferocytosis; SensTop50, sensome top50; Phago+, positive regulation of phagocytosis; MicNeu, microglia-neuron interaction; Neuroprot, neuroprotection; IFN_ISG, interferon-stimulated genes; * significantly differential gene expression (B, D: log2FC > 0.5, p-adjusted < 0.05) or GSA activity (C, F: log2FC > 0.25, p-adjusted < 0.05).

These transcriptomic changes were accompanied by morphological alterations in repopulated microglia, including increased cell and soma circularity and higher spatial complexity, indicative of a less amoeboid and more ramified morphology (Fig. S3). Together, these observations are consistent with a shift toward a more reactive and neuroprotective profile in repopulated microglia, particularly within the homeostatic compartment, relative to the vehicle group.

### Short-term microglia depletion accelerates fine motor recovery after stroke

To assess motor recovery, we chose the dMCAo model as it specifically affects fine motor function that can be evaluated using the SPR task (Fig. 3A). Paw preference was determined during a training phase and was comparably distributed across groups: stroke PLX-treated (STR PLX: 8 right, 8 left); stroke vehicle-treated (STR VEH: 8 right, 7 left); sham PLX-treated (SH PLX: 1 right, 8 left); sham vehicle-treated (SH VEH: 2 right,7 left). At the time of intervention, animals of all groups did not differ significantly in weight (SH VEH 25.34 ± 0.60 g, SH PLX 25.44 ± 0.68 g, STR VEH 25.33 ± 0.61 g, STR PLX 25.34 ± 0.49 g; one-way ANOVA, p = 0.997).

**Figure 3.**
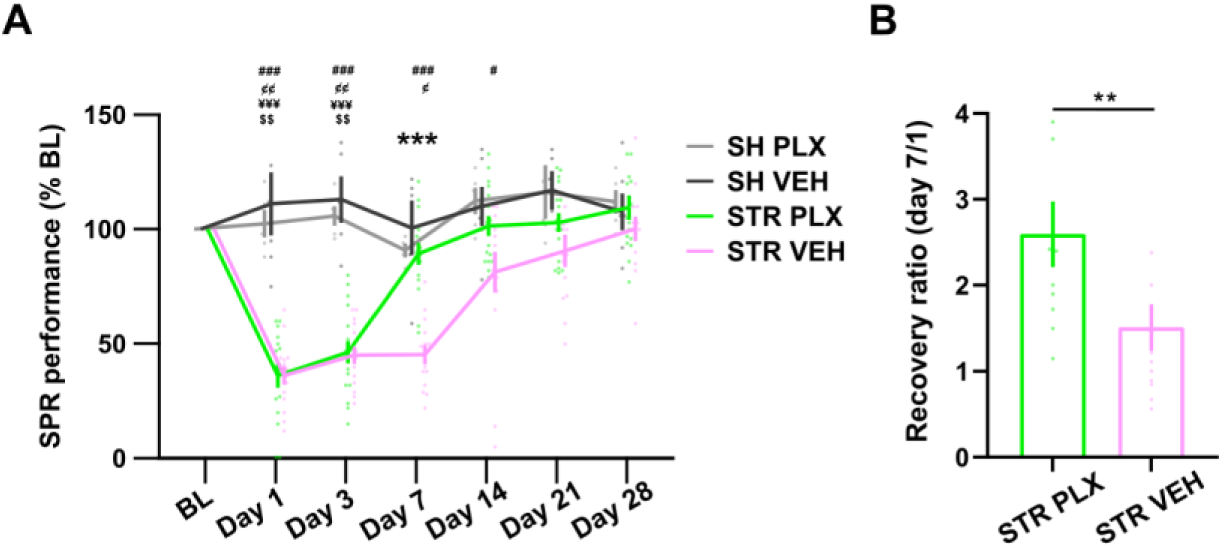
Short-term microglia depletion after stroke accelerates motor recovery in single-pellet reaching (SPR) task. (**A**) SPR performance relative to baseline (%). STR PLX_1_: n = 16, STR VEH_2_: n = 14, SH PLX_3_: n=6, SH VEH_4_: n = 5. *** p < 0.001 vs groups 1 and 2; $$ p < 0.01 vs group 1 and 3; ¥¥¥ p < 0.001 vs group 1 and 4; ¢ p < 0.05 vs group 2 and 3; ### p < 0.001 vs group 2 and 4. (**B**) Recovery ratio (day7/1) direct comparison between the stroke groups. 1 = no change; > 1 = improvement. Values above 4 were excluded for visualization; statistical analyses include all data. *, $, ¥, ¢, # p<0.05 Data are shown ± SEM. BL, baseline; SH, sham; STR, stroke; PLX, PLX5622; VEH, vehicle.

Consistent with prior work, dMCAo induced a marked drop in SPR performance on day 1 in comparison to shams (STR PLX and STR VEH vs. SH PLX: p=0.007 and p=0.009, respectively; vs. SH VEH: both p<0.001), with no differences between stroke groups (p > 0.999) ^15^. On day 7, stroke PLX-treated animals showed a drastically improved performance, nearly reaching baseline levels and no longer differing significantly from sham animals (SH PLX: p >0.814; SH VEH: p=0.994), while the stroke vehicle-treated animals were still showing significant impairments (SH PLX: p=0.026; SH VEH: p<0.001). In direct comparison, PLX-treated animals after stroke exhibited a 1.8-fold greater recovery between day 1 and day 7 than vehicle-treated mice after stroke, with median recovery rates of 2.255 vs. 1.275, p = 0.004 (Fig. 3B).

On day 14, PLX-treated stroke animals showed a sustained functional improvement (% SPR baseline) while vehicle-treated stroke animals still demonstrated slight impairments (% SPR baseline, SH VEH, p=0.043) which did not reach statistical significance in comparison to PLX-treated animals (SH PLX, p=0.139; STR PLX, p= 0.213). From day 21 onward, no differences were observed between any groups (all p ≥ 0.140).

Throughout the entire study, no differences were detected between both sham groups (day 1: p=0.934; day 3: p=0.912; day 7: p=0.851; day 14: p=0.995; day 21: p>0.999; day 28: p=0.968), confirming that PLX alone did not affect task performance in uninjured animals. Spontaneous behavior assessed in the open field did not differ significantly between the two stroke groups. Open field test showed comparable locomotor activity, exploration, and movement patterns across conditions, and no differences in weight were observed throughout the study (Fig. S4).

In summary, timed microglia depletion led to a faster recovery of fine motor function of the contralateral forepaw. Moreover, this functional improvement remained stable during the repopulation of microglia and persisted until microglia count reached normal values.

### Increased connectivity within ipsilesional M1 informs functional recovery

To investigate the underlying neurophysiological processes associated with enhanced regenerative capacity, we recorded ECoG signals from both hemispheres, covering extensive areas of the motor and sensory cortices (Fig. 4A). In our previous work, we had identified the area covered by electrode ch3, within the ipsilesional M1, as a key region involved in recovery ^15^. Given the multifaceted role of microglia in modulating synaptic turnover and receptor composition, we hypothesized that microglial depletion might result in significant alterations in functional network connectivity.

**Figure 4.**
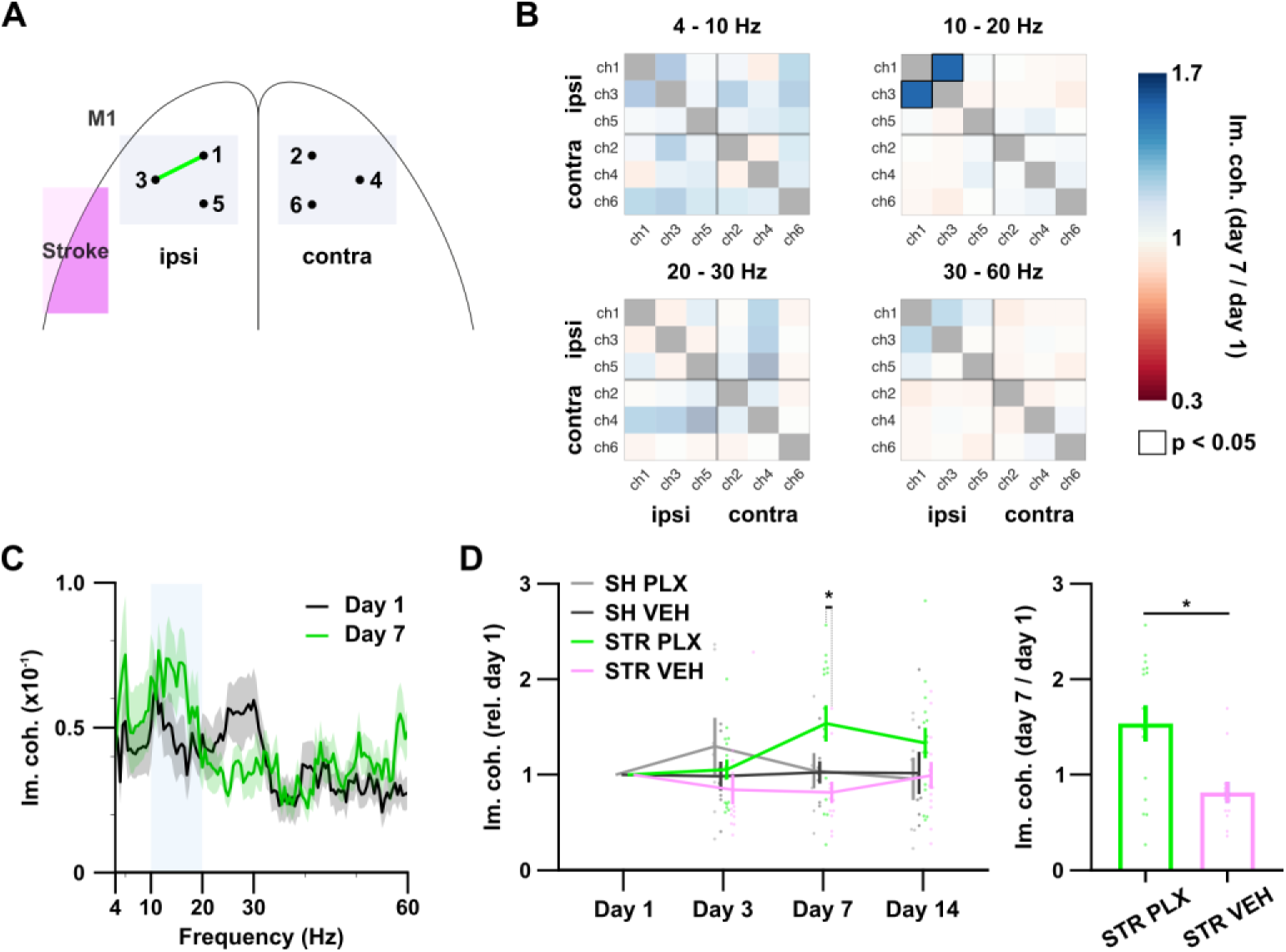
Increase of 10-20 Hz connectivity (imaginary coherence) in ipsilesional M1 of PLX-treated mice. (**A**) Schematic illustrating electrode location within M1, green line indicates significant increase of imaginary coherence (im. coh.) compared to the vehicle-treated stroke group. (**B**) Heat maps depicting increases and decreases of imaginary coherence on day 7 in relation to day 1 in given frequency bands for all possible electrode combinations (color-coded). Significant alterations compared to the vehicle-treated stroke group are outlined. (**C**) ch1-ch3 imaginary coherence spectrum of PLX-treated mice on day 1 (black) and day 7 (green) after stroke. (**D**) Left, the fold-change of ch1-ch3 10-20 Hz imaginary coherence in all four groups in relation to day 1. Right, direct comparison of the evolution imaginary coherence in the two stroke groups. SH PLX: n = 8; SH VEH: n = 7; STR PLX: n = 14; STR VEH: n = 12. * p<0.05. Data are shown ± SEM. SH, sham; STR, stroke; PLX, PLX5622; VEH, vehicle.

To assess connectivity, we calculated imaginary coherence during the recovery phase spanning from day 1 to day 7 post-stroke ^25^. For spectral and connectivity analysis, we defined frequency bands as follows: theta (4–10 Hz), low beta (10–20 Hz), high beta (20–30 Hz), and gamma (30–60 Hz).

First, we examined the connectivity between all possible electrode connections over ipsilesional and contralesional M1 during active episodes, defined by locomotion and exploratory behavior (Fig. 4A). Connectivity in the lower beta band was significantly increased, by up to ∼150%, in the post-stroke-PLX-treated group compared to vehicle-treated stroke animals. This increase was only evident between electrodes ch1 and ch3 in the ipsilesional M1 on day 7 (mean diff. 0.723, p=0.017, repeated-measures ANOVA; Fig. 4B, C, D) and was observed only during active, not resting, periods (day 7: mean diff. 0.155, p=0.969, repeated-measures ANOVA; Fig. S5D). Supporting analysis showed that the increase in beta band connectivity spanned a broadband frequency range between 10 and 20 Hz and did not coincide with the ∼16 Hz peak observed in the power spectrum (PSD mean 15.63±0.19 Hz vs. im. coh. mean 13.49±0.39 Hz, p<0.001, Welch’s t-test; Fig. 4C, Fig. S5A, B). Analysis of the aperiodic exponent of the power spectrum in M1, a marker of cortical excitation-inhibition balance, did not reveal significant differences between PLX-treated and vehicle-treated post-stroke animals (Fig. S5E). Furthermore, the observed increase in was neither correlated with spectral power in the corresponding frequency band nor with aperiodic markers (Table S4).

Next, we assessed interhemispheric connectivity between the motor and sensory network (Fig. 5A, B). In PLX-treated animals, a significant increase in theta connectivity between ipsilesional motor and sensory cortices (ch3–ch9) was observed when compared to vehicle-treated stroke animals (day 7: mean diff. 0.513, p=0.043, day 14: mean diff. 0.960, p=0.039, mixed-effects model; Fig. 5B). This effect appeared to be primarily driven by a decrease in connectivity within the vehicle-treated stroke animals over time (Fig. 5C). On the contralateral side, we found an increase in lower beta connectivity (ch4–ch14) beginning as early as day 3 (day 7: mean diff. 0.540, p=0.28, mixed-effects model; Fig. 5B, C), and a significant decrease in gamma connectivity (ch6–ch12) emerging by day 7 (STR PLX vs. STR VEH day 7: mean diff. -0.410, p=0.031, SH PLX vs. STR PLX day 7: mean diff. 0.318, p= 0.032, day 14: mean diff. 0.412, p=0.011, mixed-effects model; Fig. 5B, C). No significant changes were detected in the upper beta frequency band (Fig. 5B). While the ipsilesional increase in theta connectivity correlated with theta power in the motor cortex (ch3 4–10 Hz power vs. ch3–ch9 4–10 Hz im. coh., r=-0.434, p=0.019; repeated-measures correlation), no such relationship was found for other channel pairs (Table S4).

**Figure 5.**
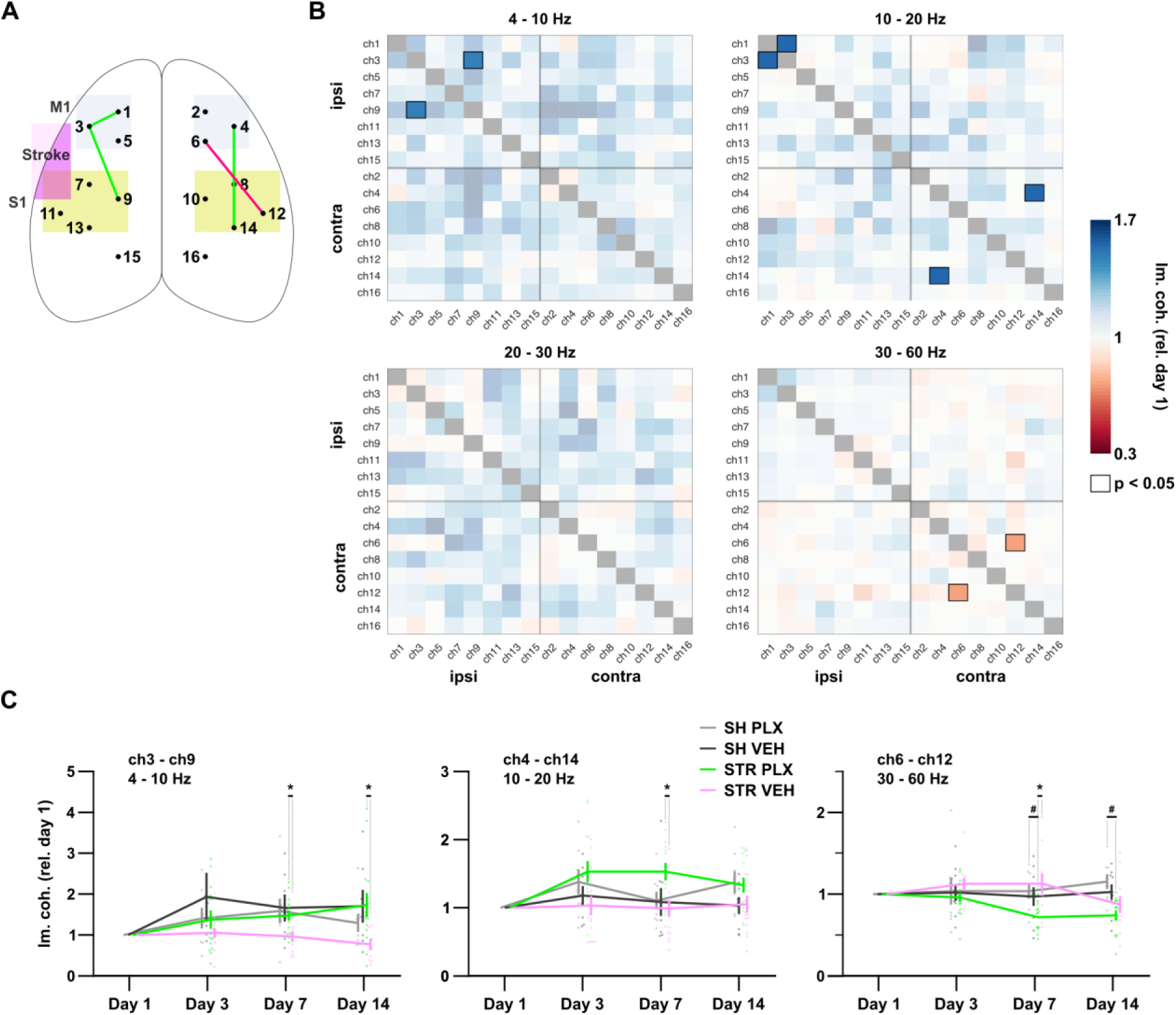
Bilateral modifications of sensorimotor networks following ischemic stroke in PLX-treated mice. (**A**) Schematic illustrating all electrode locations, green line indicates significant increase and red line significant decreases of imaginary coherence (im. coh.) compared to the vehicle-treated stroke group. (**B**) Heat maps depicting increases and decreases of imaginary coherence on day 7 in relation to day 1 in given frequency bands for all possible electrode combinations (color-coded). Significant alterations compared to the vehicle-treated stroke group are outlined. (**C**) The fold-changes of imaginary coherence in the given frequency bands and between given electrode combinations in all four groups in relation to day 1 are demonstrated. SH PLX: n ≥ 8; SH VEH: n ≥ 7; STR PLX: n ≥ 14; STR VEH: n ≥ 12; exact animal numbers varied depending on electrodes considered. *, # p<0.05. Data are shown ± SEM. SH, sham; STR, stroke; PLX, PLX5622; VEH, vehicle.

To assess whether these observations relate to improvements in fine motor skills, we examined the link between functional connectivity and motor recovery using repeated-measures correlation during the recovery phase (day 3 to day 14). M1 ipsilesional connectivity in the low beta range positively correlated with performance in the SPR task, supporting a potential mechanistic link between local network reorganization and functional recovery (r=0.379, p=0.039, repeated-measures correlation; Fig. 6B). The behavioral correlation was specific to M1 and was not found for ipsi- or contralesional sensorimotor connectivity in the theta and low beta range respectively (Fig. 6A, C). Conversely, contralesional sensorimotor gamma connectivity was significantly and inversely correlated with fine motor improvement (r=-0.415, p=0.025, repeated-measures correlation; Fig. 6D). In summary, we identified bilateral alterations in sensorimotor network dynamics following stroke. Among these, the increase in low beta-band connectivity within the ipsilesional motor cortex emerged as a strong candidate mechanism underlying the restoration of contralateral forepaw function. Additionally, reduced gamma synchronization in the contralateral sensorimotor system was likewise associated with fine motor recovery.

**Figure 6.**
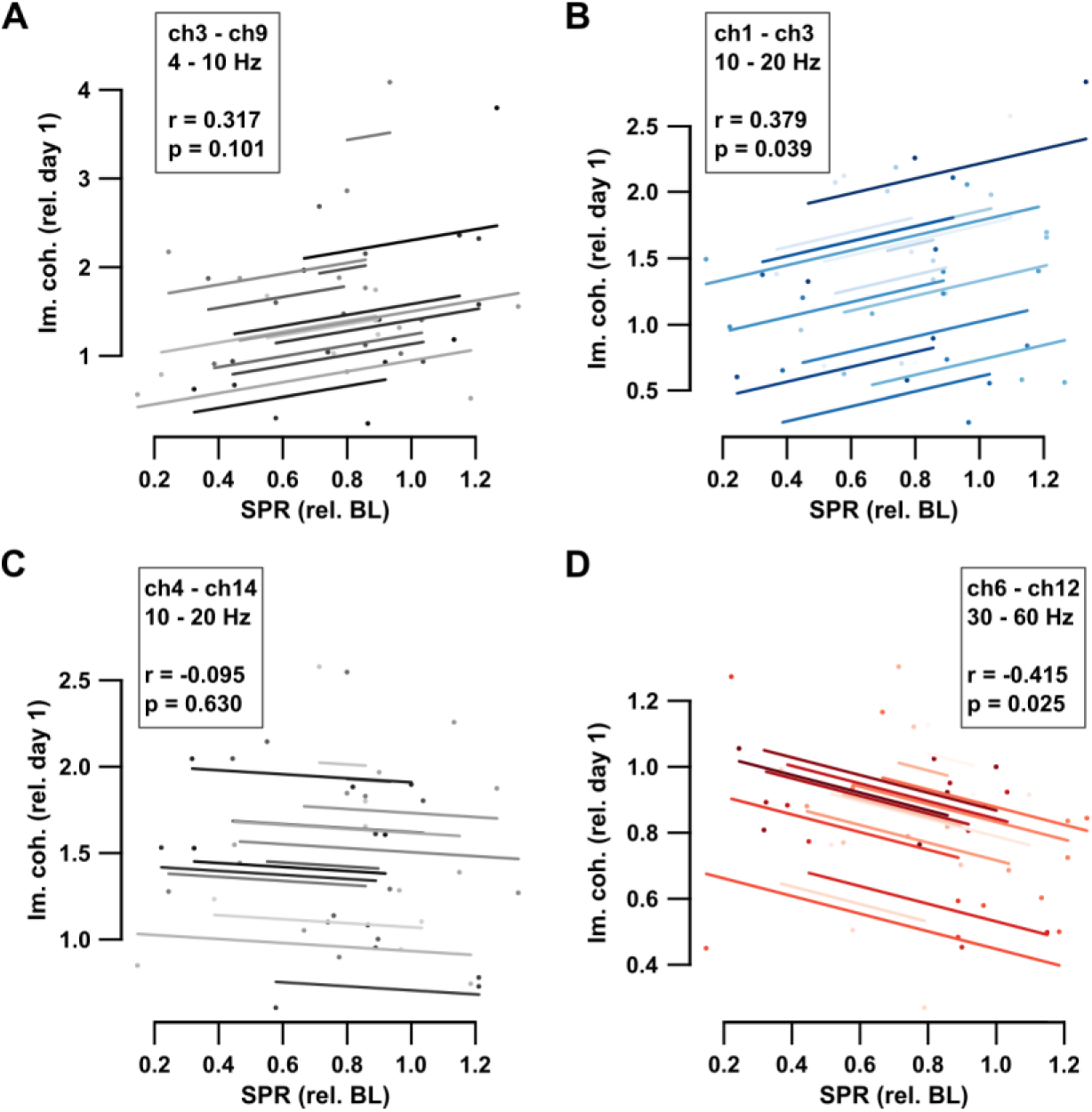
Lower beta connectivity within ipsilesional M1 of PLX-treated mice correlates with behavioral recovery. (**A**), (**B**), (**C**) and (**D**) Repeated-measures correlations of imaginary coherence (in relation to day 1) between given electrode pairs in given frequency bands and behavioral performance (in relation to baseline, BL) are shown for the recovery phase (day 3, day 7 and day 14) in PLX-treated mice after ischemic stroke. Blue lines indicate significant positive correlation, red line significant negative correlation. n = 14.

## Discussion

Here we show for the first time that short-term microglia depletion between days 3 and 7 after sensory cortical stroke in mice accelerates fine motor recovery and modulates functional connectivity within bilateral sensorimotor networks. While the depletion itself did not affect lesion size, spontaneous behavior, or general locomotion, it led to a significant early improvement in skilled forelimb use. This early recovery coincided with the peak of depletion, correlated with increased connectivity within the ipsilesional motor cortex and persisted throughout microglial repopulation, suggesting that microglial activity during this subacute phase critically shapes the trajectory of post-stroke recovery.

Microglia, as the resident immune cells of the central nervous system, are rapidly activated in response to cellular injury. In the immediate aftermath of ischemic stroke, microglia accumulate at the lesion site, where they release pro-inflammatory mediators that sustain neuroinflammation and can contribute to secondary neuronal damage ^26^. Simultaneously, they play a beneficial role by clearing necrotic tissue and secreting anti-inflammatory markers and growth factors, thereby supporting vascular remodeling and tissue repair ^27^. This dual role highlights the temporally and spatially heterogeneous impact of microglia on post-stroke recovery ^28,29^. Consequently, therapeutic strategies targeting microglial activation states or phenotypes are of increasing interest. Microglia depletion prior to ischemia consistently leads to worse outcomes, including larger infarcts and impaired functional recovery, underscoring their protective function during the acute phase ^30–33^. Similarly, sustained depletion from the acute through the chronic phase leads to detrimental behavioral outcomes ^34^. In contrast, transient microglia depletion limited to the subacute phase—particularly between days 3 and 7 post-stroke—has emerged as a promising approach ^14^. This intervention has been shown to reduce neuroinflammation, enhance motor and cognitive recovery, and shift microglial gene expression profiles toward a more homeostatic, less reactive phenotype following repopulation ^11–14^. In line with these findings, our data demonstrate that timed microglia depletion between days 3 and 7 after sensory cortical stroke leads to improved performance in the single-pellet reaching task by day 7 post-stroke, corresponding to the peak of depletion, whereas vehicle-treated controls required 14 to 21 days to reach similar functional levels. Notably, the behavioral improvement persisted through the phase of microglial repopulation, reaching day 28 post-stroke, at which point the microglial population was fully restored. These findings suggest that short-term disruption of microglial activity during a defined subacute period can initiate an accelerated and enduring recovery process. This interpretation is supported by Elmore et al.^35^, where it was demonstrated that replacing microglia improves cognitive, synaptic, and neuronal functions, suggesting that in our study activated residual microglia may inhibit recovery while newly repopulated microglia facilitate tissue repair. Consistent with this, our scRNA-seq analyses reveal that repopulated microglia exhibit transcriptional programs linked to repair and neuroprotection, in line with previous reports ^14,27^. However, the precise molecular pathways mediating these effects remain to be determined.

Microglial activation states exert a profound influence on neuronal network dynamics. Both short- and long-term microglial depletion have been shown to impact synaptic structure and function ^3,36^. Chronic depletion results in increased dendritic spine density and a greater number of synaptic contacts ^9,11,36^. In the visual cortex, microglial depletion led to a transient increase in both excitatory and inhibitory synapses on excitatory neurons, accompanied by heightened activity of both excitatory and inhibitory cell populations and these changes were reversible following repopulation ^37^. At the synaptic level, tumor necrosis factor-alpha (TNF-α) released by activated microglia modulates synaptic receptor trafficking, influencing both AMPA and GABA_A_ receptor dynamics through endocytotic and exocytotic mechanisms ^5,38^. Moreover, activated microglia have been shown to slow gamma oscillations in situ, although the underlying synaptic mechanisms remain incompletely understood ^7^. Behaviorally, microglial depletion has been associated with enhanced spatial memory and improved recovery following hippocampal injury ^9,11,36,39^. Collectively, these findings indicate that microglial depletion induces both structural and functional changes that influence cortical information processing and, ultimately, motor and cognitive performance. The rapid restoration of contralateral skilled motor function and the observed modifications of bilateral sensorimotor networks suggest that targeted suppression of activated microglia during a critical subacute window after ischemic injury may facilitate synaptic plasticity and network reorganization, thereby enhancing functional recovery.

Several human studies have investigated the relationship between cortical reorganization via functional connectivity and behavioral outcome following ischemic stroke ^40–42^. From a neuroanatomical and systems-level perspective, the observed increase in functional connectivity within the ipsilesional motor cortex in our study is particularly noteworthy, as it was significantly associated with improved contralateral motor performance. This region comprises not only the primary motor cortex (M1; channel 3) but also the transition zone between the primary and secondary motor cortices (channel 1, see also Table S4). Accordingly, the finding can be interpreted in line with human data demonstrating beneficial effects of enhanced connectivity between primary and secondary, more specifically, supplementary motor areas in stroke recovery ^43–46^. While studies in non-human primates have demonstrated the remarkable adaptability of the motor cortex following ischemic lesions through the immediate reorganization of motor maps, other work has identified specific patterns of synchrony between premotor and motor cortical areas during individual movement sequences in a reach-and-grasp task ^47–49^. In our data, the frequency ranges in which this strengthened coupling occurred corresponds to the lower beta band (10–20 Hz). Beta-band activity is classically associated with the sensorimotor cortex and basal ganglia and is typically defined as ranging from 13–30 Hz ^50,51^. However, recent studies have demonstrated the functional relevance of striatal 10 Hz oscillations for locomotion in mice, and growing evidence from rodent, macaque and human supports a subdivision into low (<20 Hz) and high (>20 Hz) beta components ^52–56^. Computational analyses suggest that low beta rhythms exhibit favorable properties, such as the ability to link neuronal ensembles that are otherwise segregated by gamma oscillations ^57^. Moreover, beta synchrony around 20 Hz between M1 and somatosensory regions has been implicated in sensorimotor integration in macaques ^58^. Disruptions in low beta connectivity, on the other hand, have been linked to impairments in other cortical processes in humans, such as working memory ^59–62^. Interestingly, the observed increase in motor cortical connectivity was not associated with changes in beta power nor with aperiodic markers previously linked to the cortical excitation–inhibition (E/I) balance ^15,63^. While an increase in aperiodic exponents at channel 3 was associated with deficits in fine motor performance in a previous study, no significant differences in this parameter were observed between the PLX-treated and vehicle-treated groups in the present dataset ^15^. This may suggest that mechanisms beyond a mere shift in cortical E/I balance contributed to the observed recovery. In addition, our data indicate a significant decrease in connectivity within a contralesional sensorimotor network, which temporally coincided with the early recovery of fine motor skills in the forepaw ipsilateral to this network. The neuroanatomical interpretation of this finding is less straightforward. Since ECoG recordings were acquired during free movement, it may be speculated that the early restoration of motor function influenced compensatory recruitment of contralateral networks or led to altered forelimb usage. Further studies will be required to elucidate this finding, as well as the specific roles of the two additional ipsilesional and contralesional network modifications described above. Taken together, we argue that the observed increase in low beta connectivity (<20 Hz) within the ipsilesional motor cortex may reflect the formation of expanded motor networks involving secondary motor areas and could represent a compensatory mechanism facilitating recovery. The results must be interpreted in light of the study’s limitations. It remains unclear whether repopulated microglia fully recapitulate the functional properties of their predecessors. Future research incorporating transcriptomic profiling and selective manipulation of microglial subsets will be necessary to delineate these pathways. Additionally, further studies should investigate the cellular interactions between microglia and neurons that facilitate the emergence of coherent and functionally integrated cell assemblies during recovery. Lastly, due to technical constraints, ECoG recordings were performed in the open field in the evening, after SPR testing in the morning; future studies should capture neural activity during the task.

Taken together, our findings underscore the critical role of microglia in shaping post-stroke recovery trajectories and highlight their potential as a therapeutic target through temporally controlled modulation of their activity.

## Supplemental material

Expanded Materials and Methods

Figures S1-S5

Tables S1-S3 (Excel files)

Tables S4-S5 (included in Supplemental PDF)

## Nonstandard Abbreviations and Acronyms

dMCAo: distal medial cerebral artery occlusion
SPR: single-pellet reaching
PLX: PLX5622

